# Unexpected complexity of everyday manual behaviors

**DOI:** 10.1101/694778

**Authors:** Yuke Yan, James M. Goodman, Dalton D. Moore, Sara A. Solla, Sliman J. Bensmaia

## Abstract

How does the brain control an effector as complex and versatile as the hand? One possibility is that the neural control of the hand is simplified by limiting the space of achievable hand postures. Indeed, hand kinematics can be largely accounted for within a small subspace of postures. This oft replicated finding has been interpreted as evidence that hand postures are confined to this subspace, and that leaving it volitionally is impossible. A prediction from this hypothesis is that measured hand movements that fall outside of this subspace reflect motor or measurement noise. To address this question, we track hand postures of human participants as they perform two distinct tasks – grasping and signing in American Sign Language. We then apply a standard dimensionality reduction technique – principal components analysis – and replicate the finding that hand movements can be largely described within a reduced subspace. However, we show that postural dimensions that fall outside of this subspace are highly structured and task dependent, suggesting that they too are under volitional control. We conclude that hand control occupies a higher dimensional space than previously considered, and propose that controlling the complexity of hand movements is well within the scope of the brain’s computational power.

## Introduction

From picking up a coffee cup to playing the piano, humans can engage in a wide range of manual behaviors, highlighting the staggering flexibility of the hand, whose 27 bones and 39 muscles give rise to more than 20 biomechanical degrees of freedom (DOF) for volitional movement^1,2^. This versatility is also mediated by a sophisticated neural system: the hand is one of the most densely innervated regions of the human body^3^ and the hand representation in sensorimotor cortex is disproportionately large^4^. Additionally, although primate motor cortex lacks a clear somatotopic organization^5^, it nonetheless seems to contain a specialized module for hand control^6^.

The complexity of the hand has called into question whether the central nervous system (CNS) can fluidly control such a complex effector^7,8^. An appealing hypothesis is that hand postures – which in theory can exist over 20 or more degrees of freedom – are reduced to a lower dimensional manifold to simplify the control problem^7–9^. Instead of spanning the space afforded by every DOF, hand control relies on a set of synergies that are combined to give rise to manual behaviors. Broadly defined, a synergy is a set of muscle activations or joint movements that are recruited collectively rather than individually^10^. This restriction of volitional movements of the hand to combinations of a small number of synergies is presumed to confer a number of computational advantages, including robustness to noise and facilitated learning of novel movements^8,9^. To date, many studies have looked for hand synergies in different movement contexts, with a mixture of evidence for and against the notion that the CNS indeed constrains hand movements to the subspace spanned by a limited basis set^9–18^.

The most compelling evidence for hand postural synergies stems from analysis of hand kinematics or of the muscle activations that drive them, which seem to occupy a lower dimensional manifold within the space spanned by the individual DOFs. Indeed, principal component analysis (PCA) of kinematics or muscle activations reveals that a small number of principal components (PCs) account for most of the variance in the movements or activations associated with a given behavior (e.g. grasping, playing piano, typing)^11,12,19–22^. The assumption underlying the interpretation of this dimensionality reduction is that high-variance principal components (PCs) – the synergies – are under volitional control whereas low-variance PCs reflect motor or measurement noise. Another possibility, however, is that the exquisite control of the hand is mediated by high dimensional sensorimotor signals, and that low-variance PCs are critical to achieving precise hand postures.

The aim of the present study is to assess whether hand movements exist in a manifold whose dimensionality is lower than its theoretical value, defined by the number of DOFs. To this end, we have human participants perform two manual behaviors – grasping and signing in American Sign Language – while we track their hand kinematics. We then assess the degree to which low-variance PCs are structured by quantifying the degree to which they carry information about the manual behavior. For example, humans and non-human primates precisely preshape their hands when grasping objects. Objects can thus be classified on the basis of the hand postures adopted while grasping them, even before contact. If low-variance PCs indeed reflect noise, they should bear no systematic relationship with the object to be grasped. However, if low-variance PCs reflect subtle but volitionally controlled adjustments of hand posture to better preshape to the object, these should also be highly object specific.

## Methods

### Hand kinematics

#### Experimental design

All procedures were approved by the Institutional Review Board (IRB) of the University of Chicago. Eight right-handed adult subjects between the ages of 21 to 40 participated in the experiment. During the experiment, subjects were instructed to perform two types of manual tasks: Grasping objects, and singing in American Sign Language (ASL) (**Figure 1A**). All eight subjects performed the grasping tasks and three subjects – who had prior knowledge of ASL – performed the signing task. All subjects performed one or both tasks with their dominant hand, the right one.

**Figure 1.**
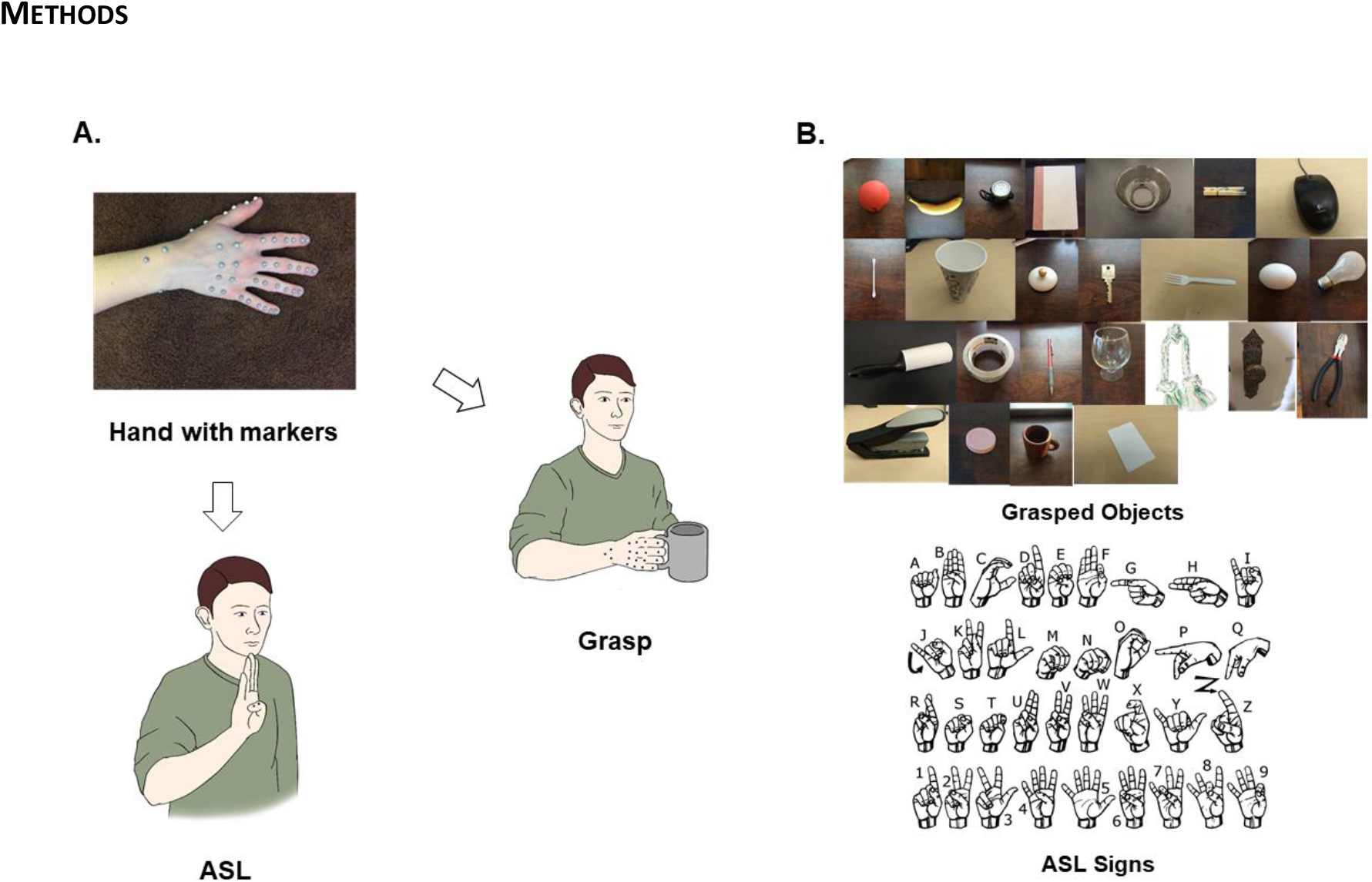
Experimental design. A| Subjects performed two manual tasks with reflective markers placed on their right hand: Grasping objects and signing in American Sign Language. Infrared cameras tracked the 3D trajectories of the markers; joint angles were calculated from them. B| Grasped objects and ASL signs.

In the grasping task, subjects began each trial by resting their right hand on a table in front of them. The experimenter then placed an object at the center of the table and subjects grasped the object with their right hand, lifted it, and held it up for approximately 1 second before replacing the object on the table and moving their hand back to the starting position. No time limit was imposed on the trial and this procedure was repeated five times for each object. Twenty five objects, varying in size, shape, and orientation, were used to elicit 30 distinct grasps (**Figure 1B**), with more grasps than objects arising from the fact that some objects could be grasped in several ways. For example, a light bulb can be grasped by the stem or by the bulb. Objects that afforded more than one grasp were presented repeatedly and subjects were cued to use a specific grasp on each presentation.

In the ASL task, subjects began at the same position as in the grasping task. On each trial, subjects signed an ASL sign – one of the 26 letters of the alphabet or a number from 1 to 10 – and repeated it five times. Again, no time limit was imposed.

#### Measurement and pre-processing

Forty one infrared-reflective markers (hemisphere shape, 4-mm diameter) were placed on the right hand of each subject, with two markers covering each finger joint, two on the ulnar, and one on the radial bone of the forearm (**Figure 1A**). Fourteen infrared cameras (8MP resolution, 250Hz) (MX-T Series, VICON, Los Angeles, CA) fixed to wall mounts and camera stands tracked the 3D trajectories of each marker, each of which was then labeled based on its respective joint using Vicon Nexus Software (VICON, Los Angeles, CA). We then calculated inverse kinematics using time-varying marker positions and a musculoskeletal model of the human arm (https://simtk.org/projects/ulb_project)^23–29^ implemented in Opensim (https://simtk.org/frs/index.php?group_id=91)^30^. The model was modified to include three rotational degrees of freedom of the first and fifth carpo-metacarpal joints, to permit reconstructions of oppositional movements of these digits. In total, we reconstructed the time-varying angles of 29 degrees of freedom, including all movement parameters of the hand and three of the wrist. We only analyzed the intervals between the start of movement and 100 ms prior to object contact or until full ASL posture.

### Analysis

#### Principal components analysis & cross-projection similarity

Kinematic synergies have been identified using principal components analysis (PCA), which expresses hand postural trajectories in terms of a set of orthogonal components, each of which reflects correlated joint trajectories. We applied PCA to the hand kinematics obtained from each individual subject^31^. To compare PC subspaces across subjects or tasks, we computed the cross-projection similarity^11^. For this, we first calculated the total variance accounted for by the first N PCs of one group (V1). Then, we projected the kinematics from the first group onto the first N PCs of a second group (V2) and calculated the total variance explained. Finally, we computed the ratio V2/V1, which approaches 1 to the extent that the second subspace resembles the first. Note that this measure is not symmetric: if the first and second groups were to change roles in an alternative V2/V1 calculation, the resulting ratio would not necessarily be equivalent. Therefore, to obtain a symmetric similarity measure, we computed the ratio V2/V1 in both directions and report the average ratio as an index of subspace similarity.

#### Classification

Next, we assessed the degree to which hand kinematics were condition specific. That is, we quantified the extent to which hand postures were dependent on the object to be grasped or the letter/number to be signed. To this end, we used linear discriminant analysis (LDA) to classify conditions based on the instantaneous hand posture (measured in joint angles) 100 ms before object contact or when the ASL sign had been achieved. Classification performance with the full kinematics provided an upper bound on the achievable classification with LDA.

Then, we gradually removed PCs in descending order of variance and projected the hand posture of each trial based on a progressively smaller subset of non-leading PCs. We then used LDA on this restricted set to classify the grasped object or the ASL posture. We used a trial-level leave-one-out cross validation: For each object, we randomly select *M*-1 trials as training data (where *M* is the total number of trials) and trained a linear discriminant classifier on the training data aggregated across objects. We then attempted to classify the combined remaining trials from all objects. We repeated this procedure *M* times (each with a different trial left out) and performance was quantified by the proportion of correct classifications.

#### Non-linear dimensionality reduction

We used two non-linear dimensionality reduction techniques to contrast with PCA: Isomap and non-linear PCA (NLPCA). We applied Isomap using the MATLAB package from Tenebaum et al^32^ with 29 nearest neighbors (though the results were robust to changes in this parameter). We calculated the variance explained by each Isomap dimension by dividing the eigenvalue of that dimension by the sum of eigenvalues. We then performed the same classification analysis by removing Isomap dimensions in descending order of eigenvalue. We applied NLPCA, an autoencoder-based approach, using the MATLAB package from Scholz et al^33^. NLPCA orders the hidden nodes (termed “nonlinear principal components”, or NLPCs) by variance explained and enforces a PCA-like structure on the low-dimensional embeddings^33^. To obtain a cumulative variance plot, we calculated the variance explained by dividing the variance of the NLPCA-reconstructed kinematics by the total variance. We performed the same classification analysis by progressively setting NLPC scores to zero prior to reconstruction of the kinematics, starting with the NLPC that accounted for the most variance and proceeding in order of decreasing explained variance.

#### Conditional Noise

One possibility is that non-isotropic, condition-dependent noise might carry information about grasped objects and support classification with low-variance PCs. To address this possibility, we denoised the kinematics, reduced their dimensionality, then added condition-dependent noise to the resulting kinematic trajectories. Specifically, we selected one trial from each object and replicated it four more times to obtain a kinematics set that contained no within-condition noise. We then reconstructed the (denoised) kinematics with only the first 10 PCs. Next, we drew from a multivariate Gaussian with zero mean and a condition-specific covariance matrix. Specifically, we randomly shuffled joint angle order and recalculated the covariance matrix of the denoised data, repeating this procedure for each object. That way, the within-object covariance (noise) was of a similar magnitude to the between-object covariance (signal) but differed systematically across objects. Finally, we scaled the conditional noise such that, when added to the 10-D data, classification performance (without removing any PCs) was similar to that from the performance with measured with the full kinematics just prior to object contact in the grasp task (~95.5%). We then performed the same classification analysis described above with sequentially removed PCs. We repeated the procedures above 5 times, each time using a different seed to generate the conditional noise (by reshuffling the joint angles and resampling from the resulting distributions).

We also examined the correlation of the scores along each PC across trials to investigate the degree to which individual PCs exhibited repeatable structure: For both the raw kinematics data and the 10-D data with conditional noise, we computed the mean correlation coefficient across trials on which the same object was presented and across trials on which different objects were presented after projecting the kinematics onto individual PCs. Note that all the noise in the simulated kinematics was condition-dependent, thereby maximizing the degree to which trial-by-trial variability in the kinematics might support classification performance.

## Results

Subjects performed two tasks – grasping various objects and signing in American Sign Language (ASL) – while we tracked their hand movements using a camera-based motion tracking system. First, we examined the degree to which the structure of hand movements was differed across tasks and individuals. Second, we ascertained the degree to which actual hand movements occupy a low-dimensional manifold of the available space spanned by the hand’s DOFs.

As might be expected, different joint trajectories were observed when subjects grasped different objects or signed different ASL letters (**Figure 2**). Moreover, the kinematics were consistent within condition – grasping a specific object or signing a specific letter – as evidenced by similar trajectories over repeated presentations of the same condition.

**Figure 2.**
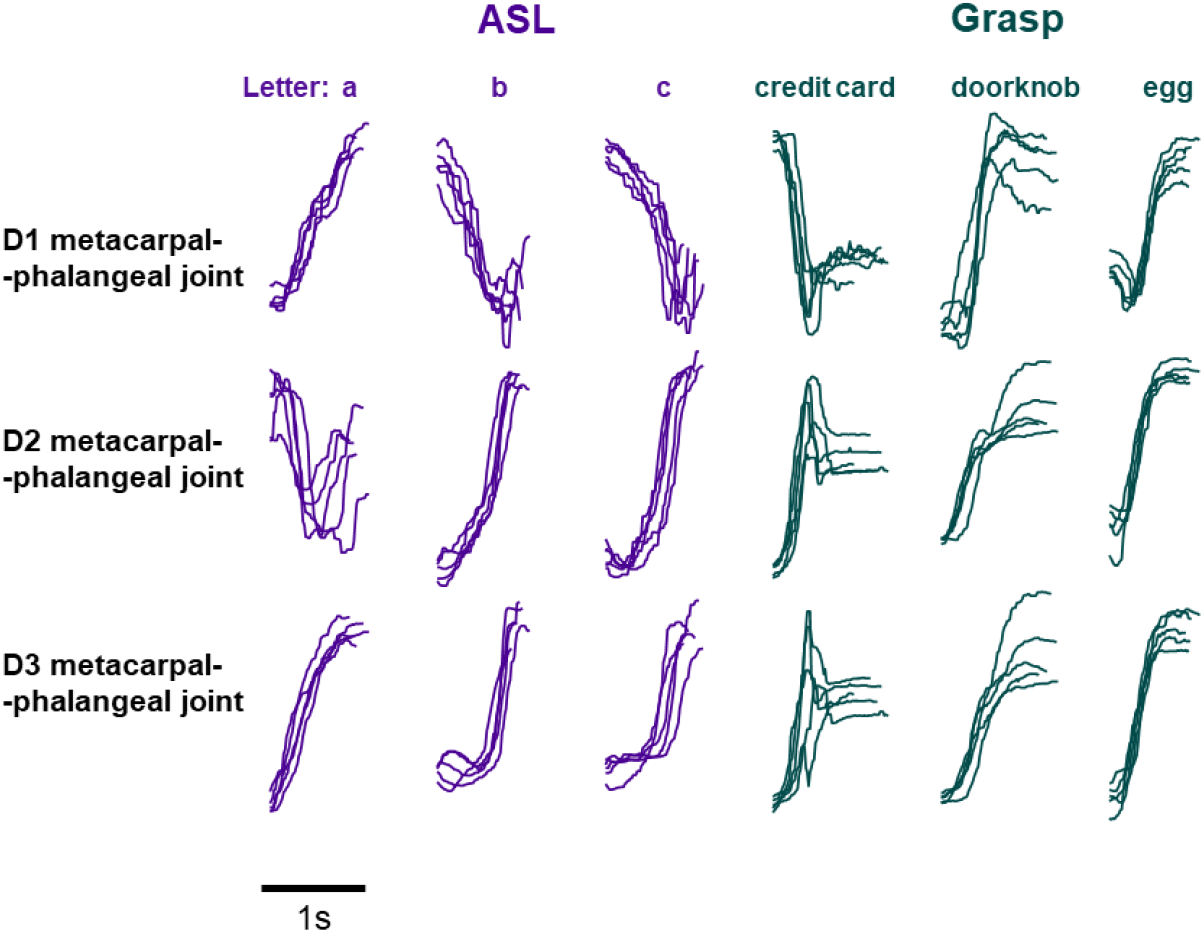
Example kinematic traces. Joint kinematics (metacarpophalangeal joints flexion of digits 1 to 3) when grasping three different objects or signing three ASL signs five times each.

### Structure of hand kinematics for two manual tasks in humans

First, we wished to reproduce previous findings that much of hand kinematics can be described within a low-dimensional subspace. To this end, we performed PCA on the joint angles and examined the cumulative variance plot. We found that 3-5 PCs were sufficient to account for 80% of the variance in the kinematics and that 8-11 principal components (PCs) accounted for 95% of variance, consistent with previous findings (**Figure 3A**)^12,34^. We then examined PCs with large and small eigenvalues (**Figure 3B**). In line with previous findings, the first two PCs of grasp and ASL involved opening and closing the hand, engaging mostly metacarpophalangeal (MCP) joint flexion/extension, some proximal and distal interphaleangeal joint flexion/extension (PIP and DIP), and some wrist flexion. One might expect PCs with small eigenvalues to reflect motor or measurement noise, and thus be unstructured. Instead, examination of low-variance PCs (the 20^th^, e.g.) revealed coordinated joint movements (e.g. ring PIP and MCP flexion in ASL) that were systematically dependent on condition (**Figure 3B**).

**Figure 3.**
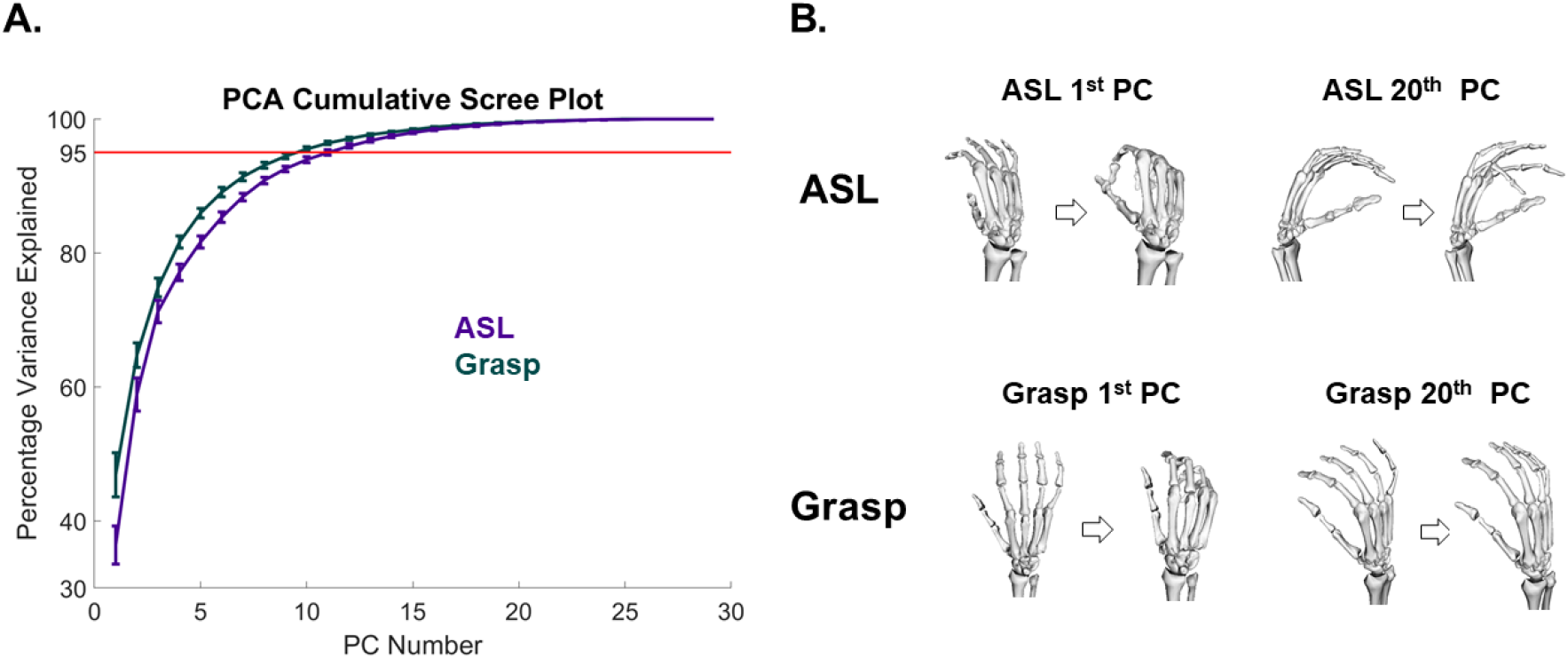
Principal components analysis. A| Cumulative variance plot showing the total percentage of variance explained vs. the number of principal components. PCA is performed separately for each task and subject. The curves are averaged across 8 subjects for grasp and across 3 subjects for ASL. B| Visualization of example PCs. For each task, we show the 1^st^ and the 20^th^ PCs (in terms of variance explained). The 20^th^ PC accounts for less than one percent of the variance.

We then compared the hand kinematics in the two tasks – grasping and ASL – by comparing their respective kinematics subspaces using cross-projection similarity. For each subject doing both tasks (*N* = 3), we calculated how much variance in the kinematics of one task were accounted for by the dimensions of the other (within-subject similarity). Despite the apparent dissimilarity of grasping and signing movements, we found that the underlying subspaces were very similar: PCs from grasping explained about 85% of the variance in ASL kinematics and vice versa (**Supplementary Figure *1***). We also found that different subjects performing the same task (grasping or signing) yielded similar subspaces (**Supplementary Figure *1***).

### Structure of low-variance PCs

Having replicated previous results that hand movements can be reconstructed with high precision using a reduced basis set, and having showed that similar basis sets accounted for movements across both tasks and subjects, we then examined whether low-variance PCs outside these basis sets were structured in a condition-specific manner. We found that the kinematics projected on the low-variance PCs across repeated presentations of the same condition (same grasped object, same ASL letter) varied systematically with the grasped object or signed letter (**Figure 4 A, B**), although this structure was noisier than that for the high-variance PCs (**Figure 4 C, D**), as might be expected. In other words, even the low-variance PCs reflect structure rather than noise in the kinematics. The trajectories along all PCs were far more consistent than would be expected by chance (**Supplementary Figure 2**). To quantify the degree to which kinematics differed across conditions, we classified objects or letters using progressively reduced kinematic subspaces that captured monotonically less variance. We found that classification accuracy was above chance even after most PCs had been removed, and that high performance was achieved even with PCs which collectively accounted for less than one percent of the variance in kinematics (**Figure 5 A, B**). A similar result was obtained when progressively removing LDA dimensions rather than PCs (**Supplementary Figure 3**). Results from these classification analyses are thus inconsistent with the hypothesis that low-variance PCs reflect motor or measurement noise. Rather, these PCs seem to reflect subtle dimensions of movement that are under volitional control and contribute to the exquisitely precise pre-shaping of the hand to an object or to the detailed execution of a complex hand conformation required to produce an ASL sign.

**Figure 4.**
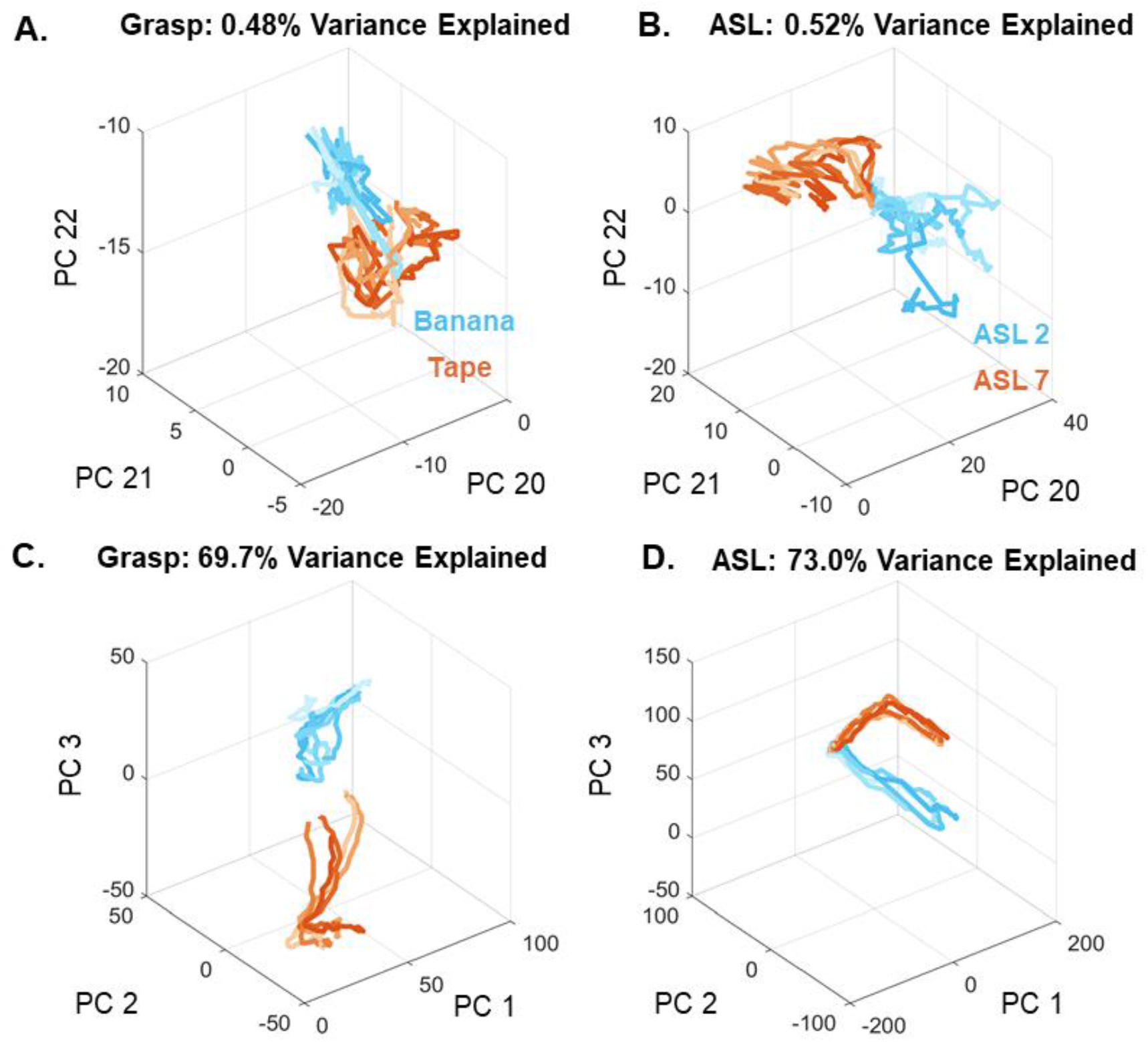
Object-specific kinematics in PC subspaces. A| Projection of the kinematics onto PCs 21 through 23 for five grasps each of tape and banana. B| Projection of the kinematics onto PCs 21 through 23 for five repeated signs each of “two” and “seven” in ASL. C| Projection of the kinematics onto PCs 1 through 3 for the conditions shown in panel A. D | Projection of the kinematics onto PCs 1 through 3 for the conditions shown in panel B. In all panels, separate objects and signs are indicated by color, and different trials are indicated by traces of different lightness.

**Figure 5.**
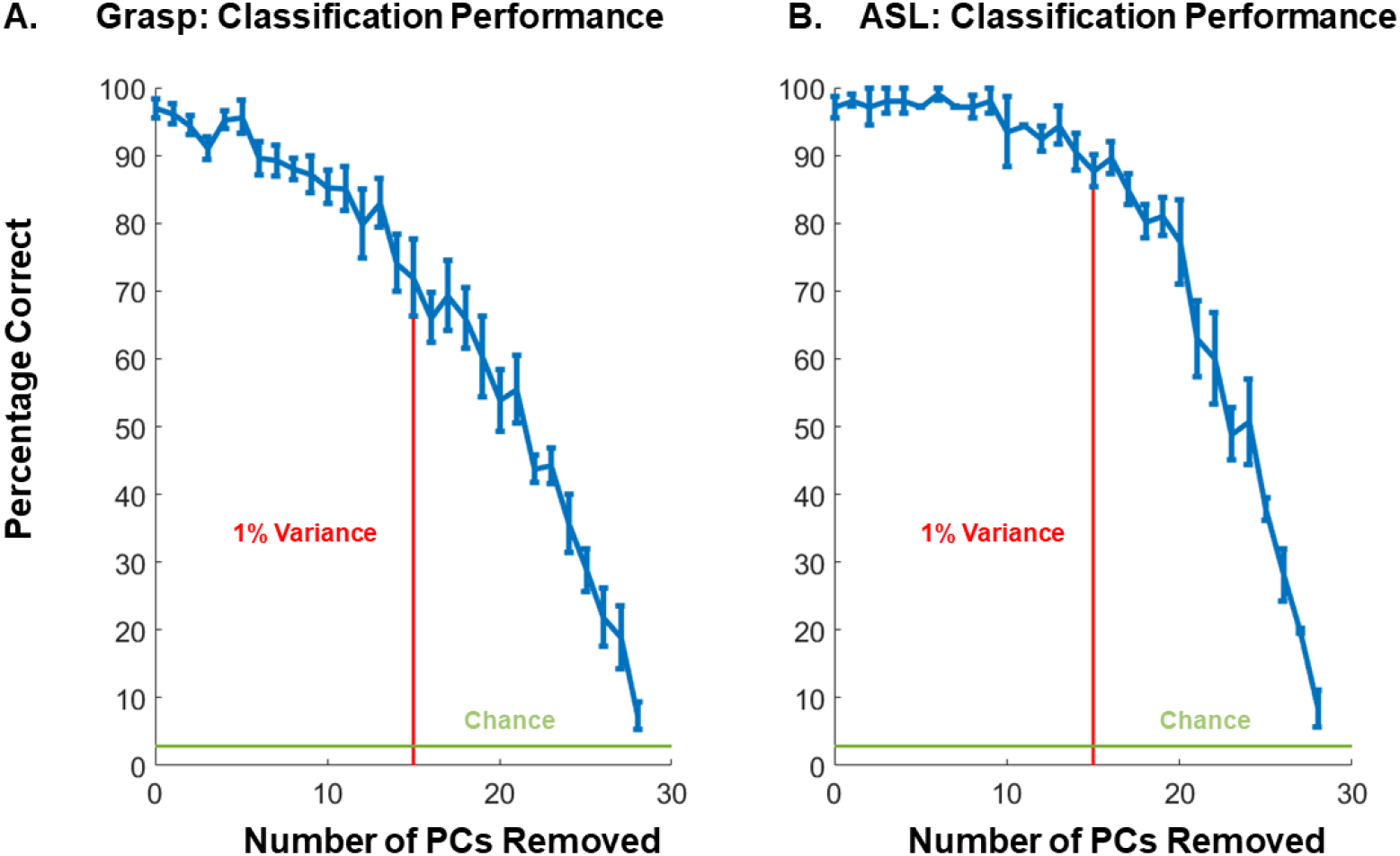
Classification of condition (object, sign) based on kinematics in progressively reduced subspaces. A| Grasp classification performance after progressively removing PCs, from high variance to low. B| ASL classification performance based on reduced kinematic subspaces. Vertical lines denote the PC beyond which each PC accounts for less than 1% of the variance. Note that objects and letters can be classified accurately based on just a handful of high-variance PCs (**Supplementary Figure 4**).

### Non-linear manifolds

We showed that low-variance PCs are structured and task-related. However, since PCA is a linear dimensionality reduction technique, w4e considered the possibility that hand kinematics occupy a low-dimensional non-linear manifold, and that the low-variance PCs reflect a linear approximation of non-linear dimensions. Indeed, such a nonlinearity could in principle explain why low-variance dimensions carry condition-specific information, thereby supporting the classification of object identity or ASL sign (**Figure 5**). To address this possibility, we performed the same classification analysis with two non-linear dimension reduction techniques, namely Isomap^32^ and non-linear PCA (NLPCA)^33^. Consistent with previous findings^35^, we found that PCA provides the most parsimonious representation of the kinematics, as gauged by variance explained (**Figure 6A**), a counterintuitive result given that PCA is restricted to linear transformations whereas the other two approaches are not. More importantly, all three algorithms yielded high classification performance (> 50 %) after removing the 20 leading dimensions, which account for more than 90% of the variance (**Figure 6B**). If hand kinematics were low-dimensional and non-linear, non-linear dimensionality reduction would account for variance more parsimoniously and a smaller number of dimensions would contain all the task-relevant information, but this is not what we found. We conclude the high information content in low-variance PCs is not a trivial artifact of non-linearity, at least of a non-linearity that could be captured by the two well established approaches to non-linear dimensionality reduction used here.

**Figure 6.**
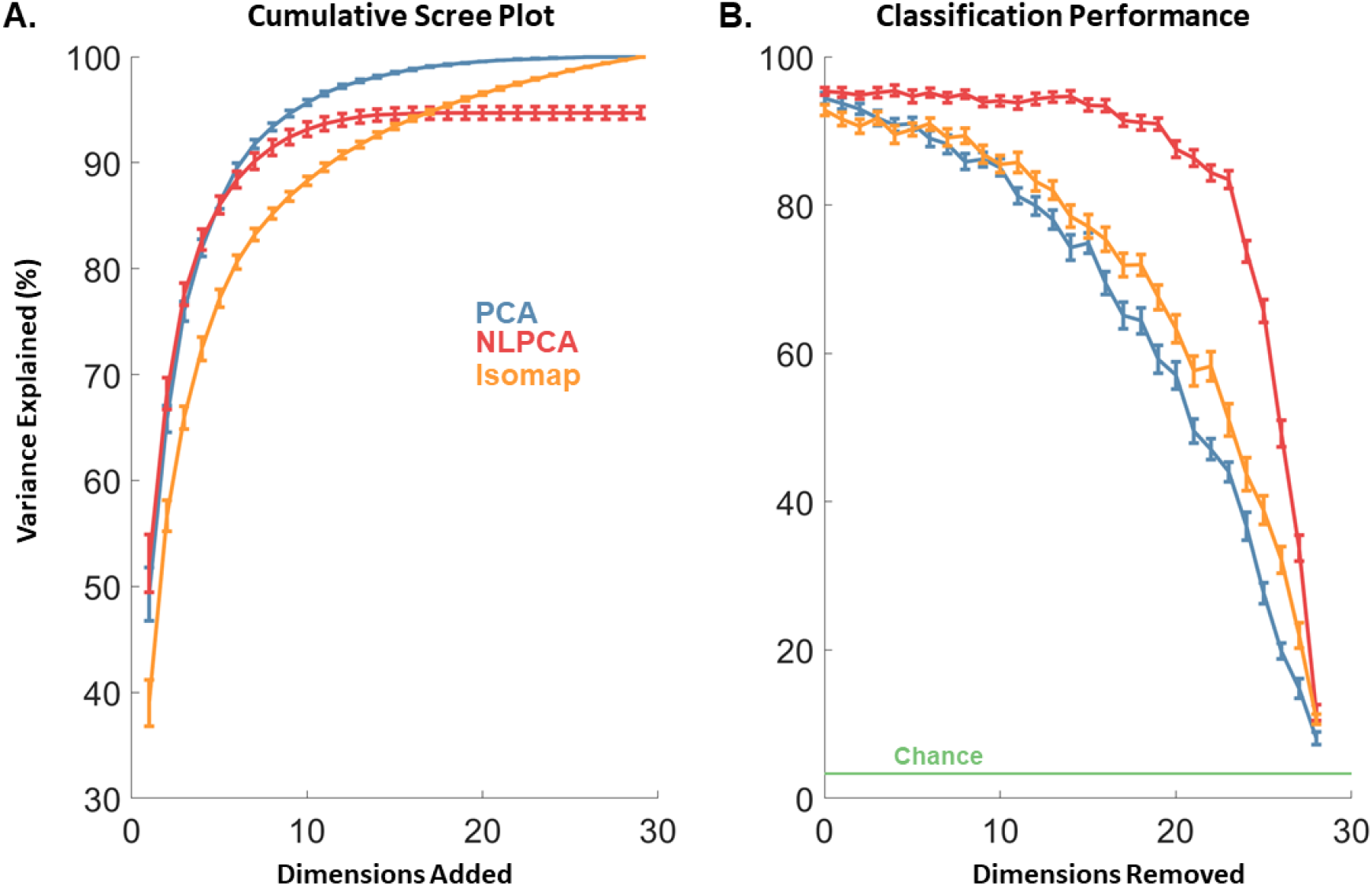
Scree plot for linear and non-linear dimensionality reduction approaches and resulting classification performance for grasp. A| Cumulative variance explained by each dimension of PCA, NLPCA, and Isomap. B| Grasp classification performance after progressively removing PCs from high variance to low, for each of the three dimensionality reduction techniques.

### Condition-dependent noise

Next, we examined whether our ability to classify objects based on low-variance PCs might be an artifact of condition-dependent noise. Indeed, one possibility is that the variability in the kinematics depends on which object is being grasped, as might be predicted by optimal feedback control^36^. This condition-dependent noise might then be exploited by the classifier. To test this possibility, we reduced the kinematics to 10 dimensions and added noise whose structure depended on the object being grasped. We then assessed the degree to which we could classify objects based on the low-variance PCs (where in principle no signal should be present, only noise). We found that, with the addition of condition-dependent noise, the first 10 PCs accounted for less variance in the kinematics than did even the original kinematics, despite the fact that all the signal is confined to those dimensions (**Supplementary Figure *5***). Furthermore, classification performance with low-variance PCs (>10) was above chance, despite the lack of true signal (**Supplementary Figure 6**). However, performance with the simulated low-dimensional kinematics dropped much faster as PCs were removed than did that with the measured kinematics (**Supplementary Figure 6**). Furthermore, the correlation between within-object PC scores was near zero for the low variance PCs (**Supplementary Figure 7**). That is, low-variance PCs of the simulated kinematics, which comprised no signal and only condition-dependent noise, were not nearly as structured as were the low-variance PCs in the measured kinematics. In summary, condition-dependent noise yields classification performance with low variance PCs that is only slightly above chance and cannot account for the observed structure in the low-variance PCs of grasping and ASL kinematics.

## Discussion

### The similarity of kinematic subspaces underlying different manual behaviors

We found that the structure of the hand postures adopted in the two tasks – grasping and signing ASL – were virtually indistinguishable. Indeed, Grasping and ASL – which each included ~30 distinct conditions – yielded subspaces that were no more different than were the kinematic subspaces of different subjects performing the same task (**Supplementary Figure 1**). At first glance, this finding seems to be inconsistent with previous observations that kinematics are more similar within task and across subjects than across tasks within subject^11,37^. However, our analysis differs from its predecessors in two fundamental ways. First, we sampled the two tasks across many conditions (objects, signs) in contrast to the more restricted conditions – such as manipulating a credit card or flipping book pages – used in previous studies. The subspaces we computed thus individually reflect a greater breadth of possible hand conformations than would subspaces computed from much more limited tasks. Second, the two tasks that we implemented did not entail contact with objects, which introduces hand conformations that cannot be achieved without contact (for example, extension of the distal interphalangeal joint). The resulting manifold thus reflects not just volitionally achieved kinematics but also the structure of the object. Systematic examination of other manual behaviors with and without object contact will be necessary to conclusively establish the degree to which contact shapes kinematics.

### Volitional hand movements are high dimensional

One hypothesis derived from (but not necessarily entailed by) optimal feedback control theory stipulates that the CNS defines low-dimensional manifolds of control to satisfy movement goals on a task-by-task basis^38^, with motor noise being preferentially shunted into dimensions outside of such manifolds^36,37,39^. According to this hypothesis, the readout of grip type or ASL letter should be restricted to a lower-dimensional manifold than that afforded by the biomechanics of the hand. Classification based on subsets of principal components revealed that low-variance (e.g., 20-22) PCs contain substantial condition specific information for a given task. Indeed, kinematic trajectories projected on these PCs were highly consistent within condition and different across conditions (**Supplementary Figure 2**). Such task-relevant information would not be present if such dimensions simply reflected motor or measurement noise, suggesting instead that volitionally controlled hand movements occupy a high dimensional space.

One might argue that non-linear dimensionality reduction is better suited to reveal the dimensionality of hand postures. However, linear approaches, including PCA, have been previously shown to yield more efficient and reliable manifolds for kinematics than do non-linear ones^35^. Not only do we replicate this result, but we also show that non-linear dimensionality reduction does not capture behaviorally relevant aspects of the hand kinematics more efficiently than does PCA (**Figure 6**). Our results are thus consistent with the hypothesis that hand postures occupy a high-dimensional manifold, even for an everyday manual behavior such as grasping.

### Implications for the interpretation of synergies

High-dimensional kinematics do not imply wholly unconstrained control of the hand. Indeed, single-finger movements attempted by monkeys^40^ and humans^41^ are never perfectly individuated: they comprise incidental movements of the other digits arising in part from co-contraction of musculature associated with other digits. Recently, the analysis of the constraints imposed on neural activity in primary motor cortex (M1) revealed that activity patterns are constrained not only to a particular linear subspace^42^, but also to a particular bounded region – a “repertoire” – within that subspace^43^. The known constraints imposed on volitional hand movements might be better explained in terms of a bounded repertoire within a high-dimensional subspace rather than an unbounded repertoire within a low-dimensional subspace.

Whether the high-dimensional structure of hand movements argues against the notion of low-dimensional motor control in general remains to be seen. Indeed, numerous reports suggest that kinematics of the hand are especially high-dimensional relative to kinematics of other effectors^16,44–46^. Moreover, a subdivision of primate M1 appears to have direct access to muscles, particularly muscles of the hand^6^, which might constitute anatomical evidence for the special nature of hand motor control. A common argument in support of low-dimensional motor control is that the brain needs to simplify the problem of controlling a complex effector such as a hand to solve it. Note, however, that control signals required for a 29 DOF-effector can occupy a neural manifold whose dimensionality is much lower than that of the possible neural space spanned by the activity of all neurons modulated by the task^47^. Recent advances in large scale neuronal recordings suggest that high-dimensional representations are possible if not common: sensory representations of natural scenes in primary visual cortex exceed 500 dimensions^48^, an order of magnitude more than the implied representations of hand postures. Nonetheless, visual percepts are highly intuitive and allow for the accurate, rapid, and effortless identification of complex objects^49^. In comparison, motor control is positively straightforward!

## Acknowledgments

We would like to thank Matt Kaufman for help with the title. This work was supported by NINDS grant R01 NS082865.

## Supplementary Figures

**Supplementary Figure 1.**
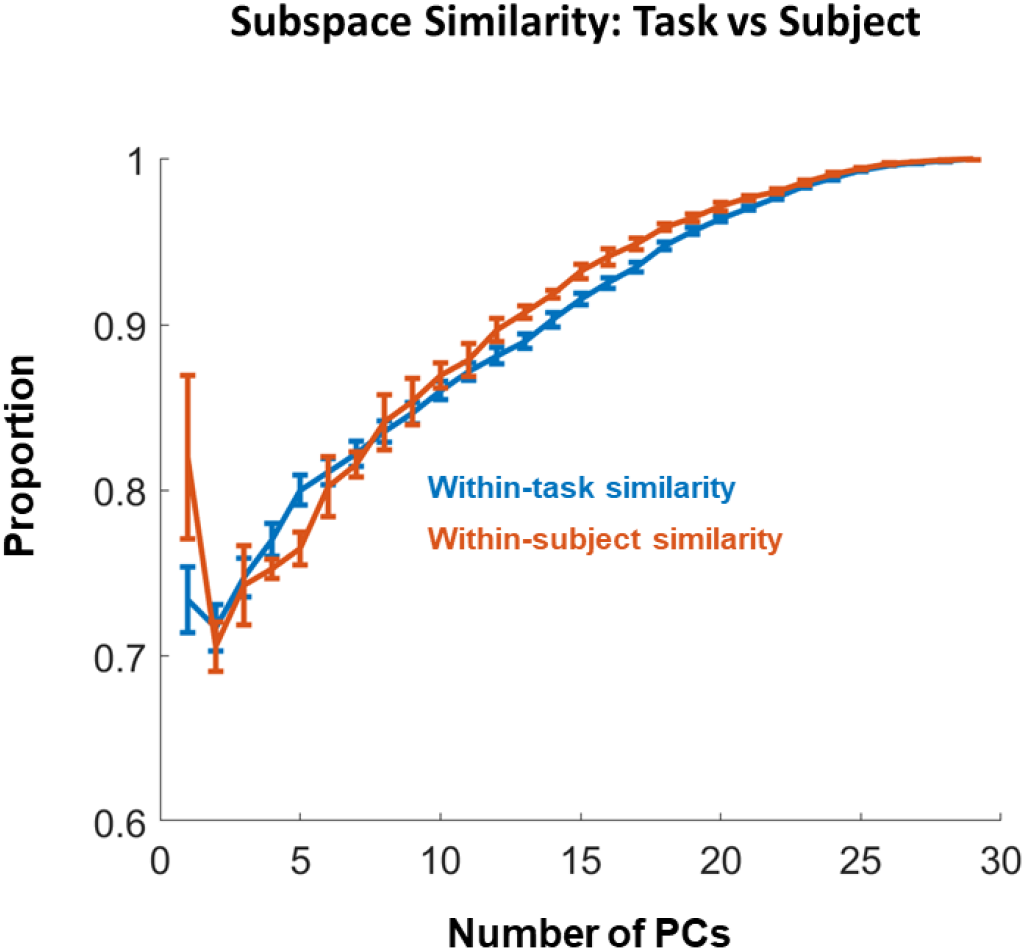
Cross-projection similarity between kinematics subspaces (see Methods for more details) within and across tasks. Briefly, cross-projection similarity refers to how much variance from one dataset can be explained by the dimensions (PCs) of the other. Within-task similarity compares the kinematics from different subjects performing the same task and within-subject similarity compares kinematics from the same subjects doing tasks (grasping and ASL). Note the similarity in the first PC for within-task similarity.

**Supplementary Figure 2.**
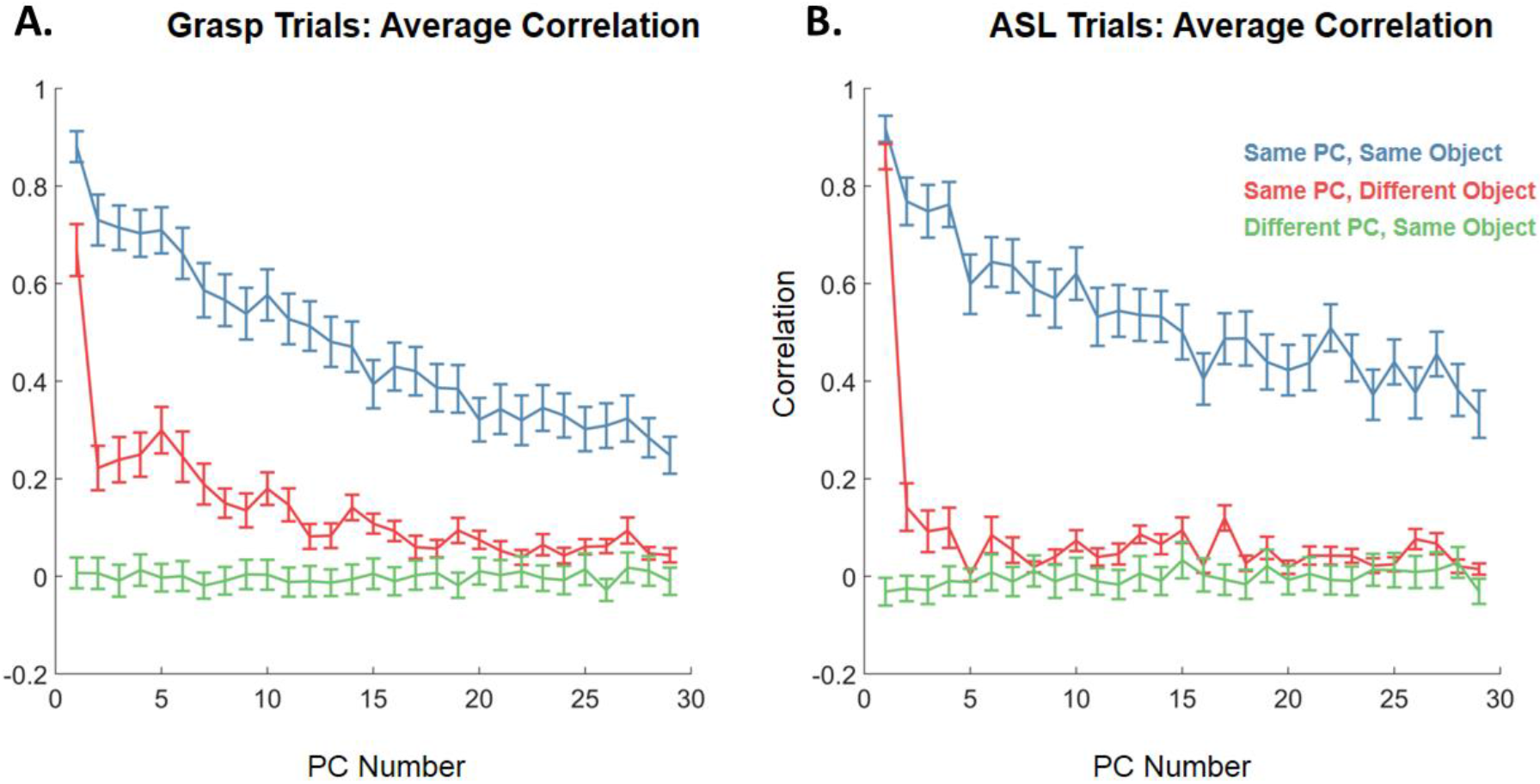
Consistency of kinematics projected onto each PC. A| Mean correlation coefficient between kinematics for different grasp trials with the same object projected onto the same PC (blue), for different grasp trials with the same object projected onto different PCs (green), and for grasp trials with different objects projected onto the same PCs (red). The within-object, within-PC correlations are systematically higher than their shuffled counterparts. B| Same analysis for ASL trials.

**Supplementary Figure 3.**
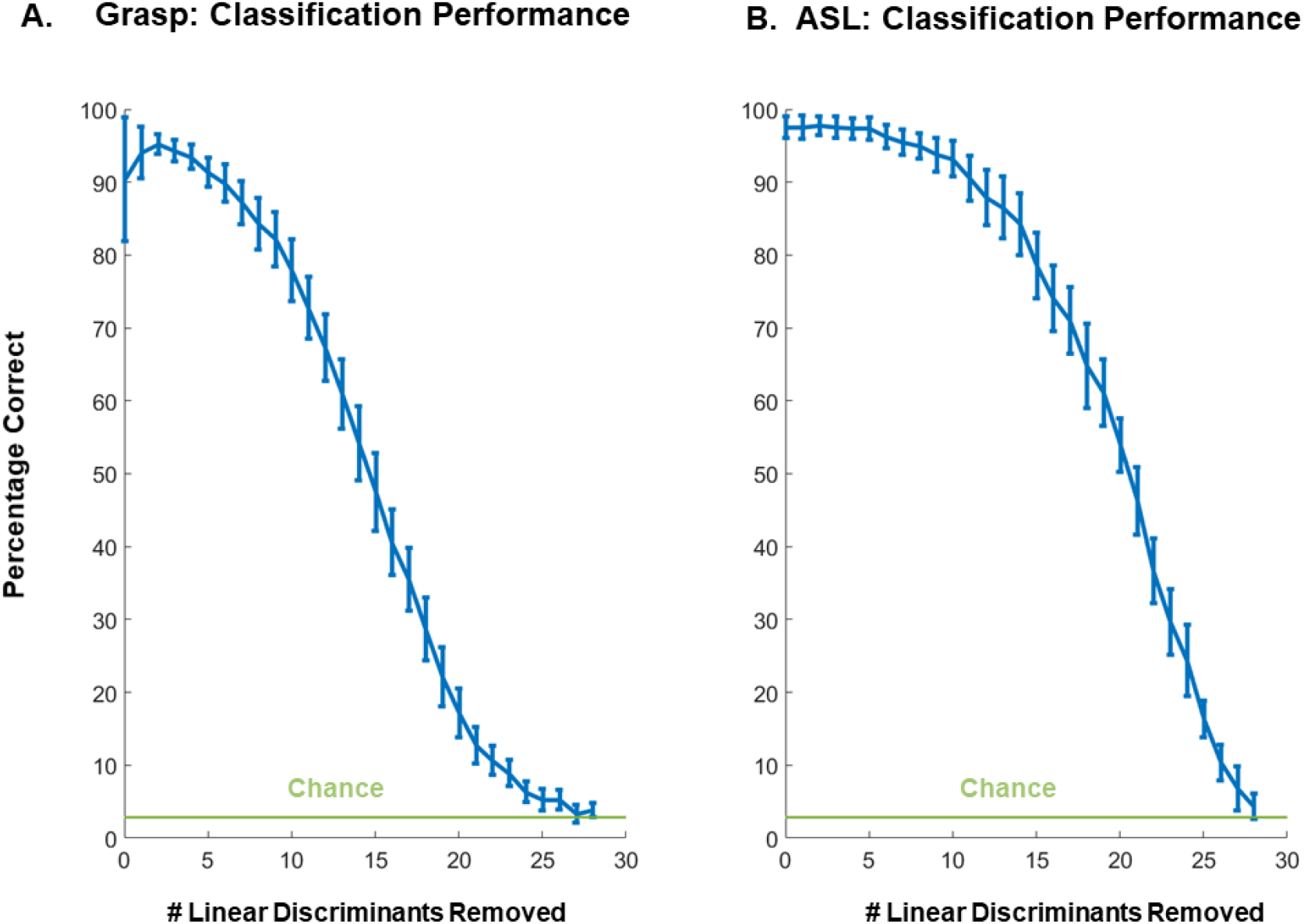
Classification performance based on a progressively reduced number of linear discriminants. A| Classification accuracy for grasped objects as the dimensions identified by LDA are removed in decreasing order of Fisher’s coefficient (ratio of between class variance to within class variance). B| Classification accuracy of ASL posture as LDA dimensions are removed in the same way.

**Supplementary Figure 4.**
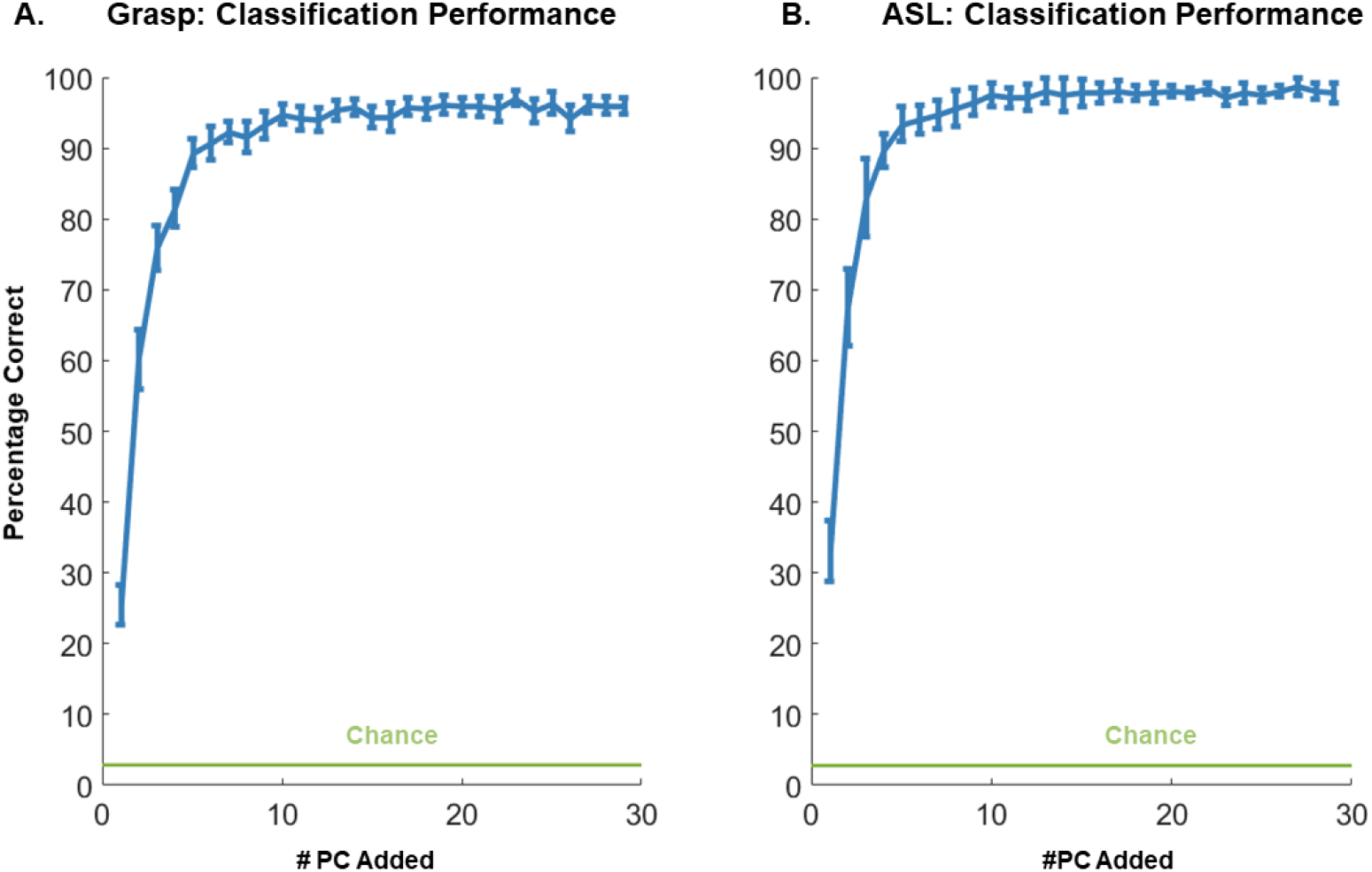
Classification performance with progressively larger subspaces. A| Classification performance for grasped objects as PCs are added in descending order of variance explained (blue curve). B| Classification performance for ASL letters vs. number of PCs. Green line denotes chance performance.

**Supplementary Figure 5.**
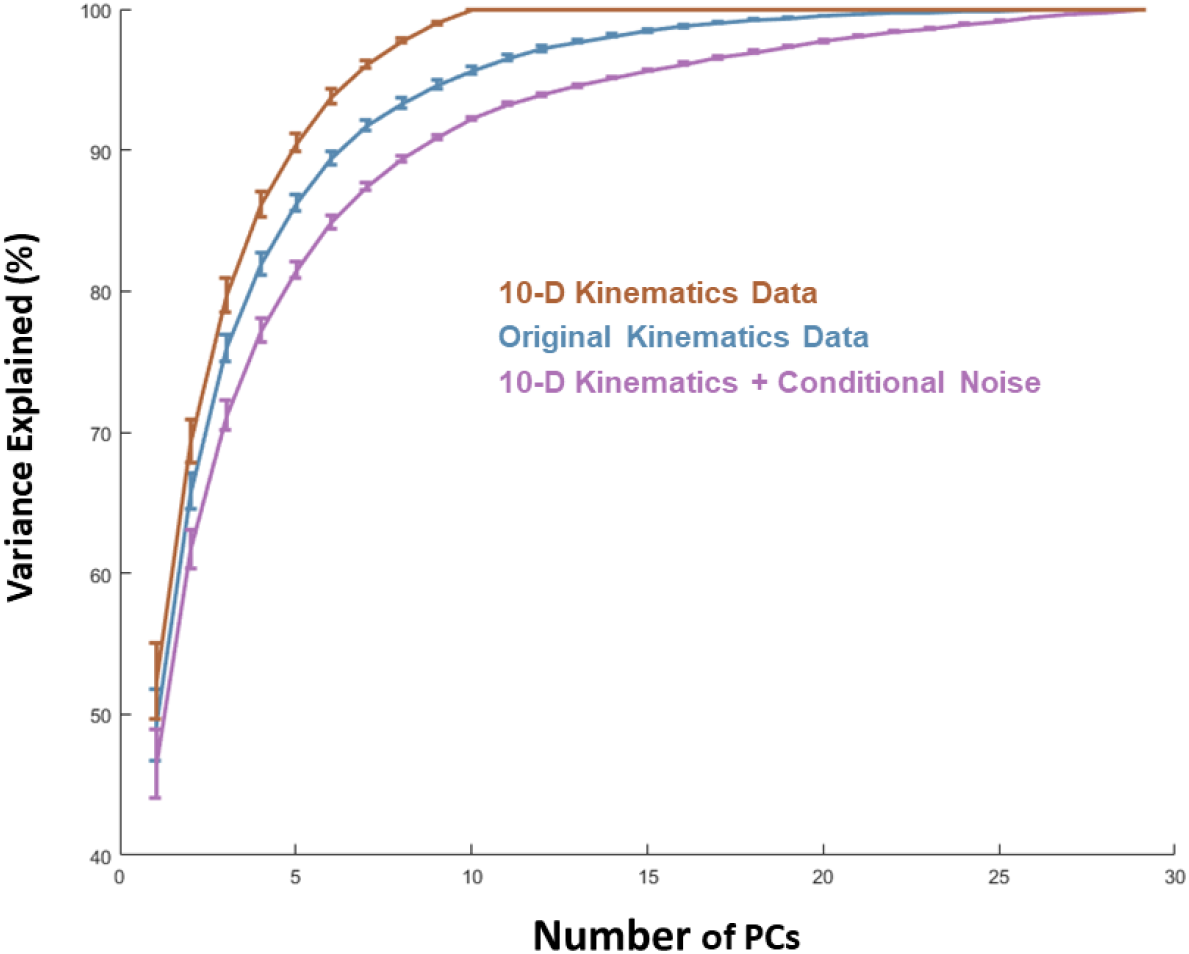
Cumulative scree plot for conditional noise simulation. Cumulative percentage of variance explained as a function of the number of PCs for the original kinematics data, 10-D denoised kinematics, and 10-D denoised kinematics + conditional noise.

**Supplementary Figure 6.**
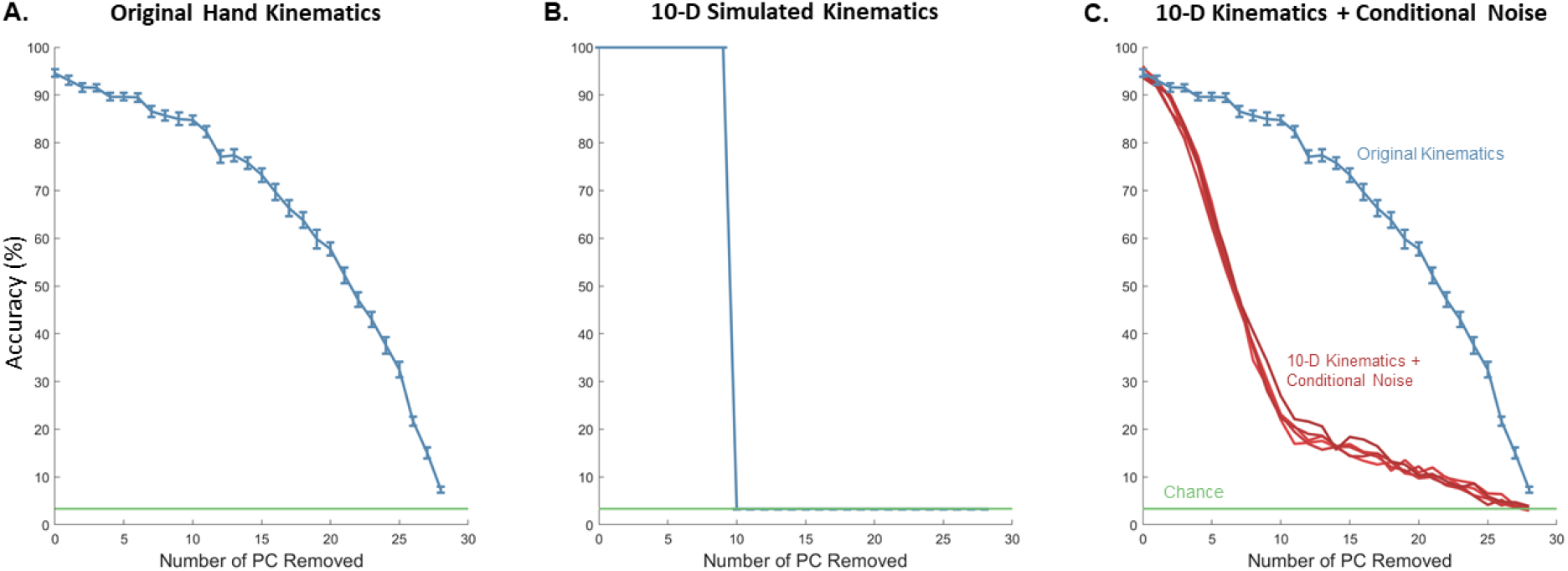
Classification with simulated low-dimensional kinematics and condition-specific noise. A| Mean classification performance of grasped objects based on the original kinematics. B| Classification performance based on 10-dimensional de-noised kinematics. In brief, the data is denoised by taking 1 trial for each object and replicating the trial for 5 times (thereby removing all variability across trials and yielding perfect classification performance through the first 10 PCs). C| Classification performance based on the simulated (denoised, low-dimensional) kinematics to which conditional noise has been added (red curves). Each red curve represents the classification results with a conditional noise generated using a different seed (5 repetitions in total). Error bars are too small to be visible. We include the result using the original hand kinematics (blue curve, same as panel A) for comparison.

**Supplementary Figure 7.**
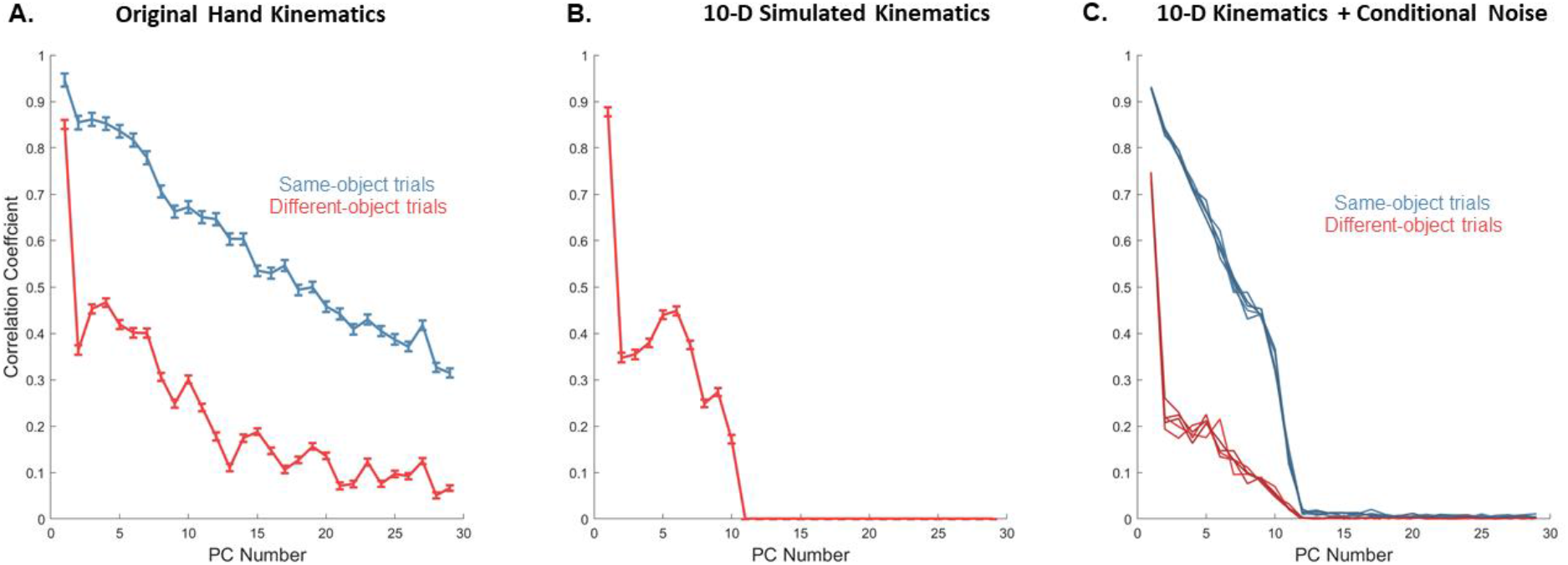
Structure in the simulated kinematics. A| Mean correlation coefficient for pairs of same-object grasp trials and different-object grasp trials projected onto individual PCs for the measured kinematics. B| Mean correlation coefficients for denoised kinematics reconstructed using only 10 PCs. Here, the same-object correlations are not shown because all same-objects trials are identical given our denoising procedures. C| Correlation coefficients for the simulated low-dimensional kinematics with added condition-specific noise. Blue curves: same object trials. Red curves: different object trials. Each red/blue curve represents the mean correlation coefficient with conditional noise simulation generated using a different seed (5 repetitions in total).

